# An actor-model framework for visual sensory encoding

**DOI:** 10.1101/2023.08.14.553166

**Authors:** Franklin Leong, Babak Rahmani, Demetri Psaltis, Christophe Moser, Diego Ghezzi

## Abstract

A fundamental challenge in neuroengineering is determining a proper input to a sensory system that yields the desired functional output. In neuroprosthetics, this process is known as sensory encoding, and it holds a crucial role in prosthetic devices restoring sensory perception in individuals with disabilities. For example, in visual prostheses, one key aspect of image encoding is to down-sample the images captured by a camera to a size matching the number of inputs and resolution of the prosthesis. Here, we show that down-sampling an image using the inherent computation of the retinal network yields better performance compared to a learning-free down-sampling encoding. We validated a learning-based approach (actor-model framework) that exploits the signal transformation from photoreceptors to retinal ganglion cells measured in explanted retinas. The actor-model framework generates down-sampled images eliciting a neuronal response in-silico and ex-vivo with higher neuronal reliability to the one produced by original images compared to a learning-free approach (i.e. pixel averaging). In addition, the actor-model learned that contrast is a crucial feature for effective down-sampling. This methodological approach could serve as a template for future image encoding strategies. Ultimately, it can be exploited to improve encoding strategies in visual prostheses or other sensory prostheses such as cochlear or limb.

## INTRODUCTION

In a biological system, sensory organs capture information from the environment and convert it into neural signals that can be interpreted and used by the nervous system. This transformation is known as sensory encoding. Similarly, an artificial system converts information from sensors into stimulation parameters of sensory prostheses. One notable example is auditory encoding in cochlear implants, where the electrical stimulation is applied in different frequency regions of the auditory nerve^1–3^. This process allows deaf individuals to regain hearing sound. Likewise, limb prostheses provide users with tactile feedback to improve their quality of life^4–11^. Similar to auditory and tactile encoding, image encoding also plays a huge role in visual prostheses. Image encoding is critical to improve the patient perception but it is not straightforward. The challenge is to find the best method to encode visual information to the resolution of a retinal^12,13^, optic nerve^14,15^, or cortical prosthesis^16,17^. Information flows from approximately 120 million photoreceptors to roughly 1.2 million retinal ganglion cells (RGCs) divided into several functional classes, which project to the lateral geniculate nucleus and then to the visual areas where further image processing occurs^18–21^.

Understanding and mimicking the principles of biological sensory encoding is imperative to develop effective prosthetic devices that can provide meaningful sensory experience to users. For example, it was recently demonstrated that a bio-mimetic stimulation elicited a more natural tactile perception^22^. In the same study, Valle et al. showed that a more natural sensory feedback increases the embodiment of the device. Further research is necessary to optimize sensory encoding methods for prosthetic devices^23^. In essence, sensory encoding in neuroprosthetics is a form of dimensionality reduction. High-dimensional information from sensors is encoded into stimulation parameters of implanted electrodes. However, the number of electrodes in prostheses is usually much fewer than the number of biological cells in the sensory system^24,25^. Hence, finding a proper artificial input that can elicit the desired perception is an ill-posed problem: there are multiple inputs that could possibly yield the same output. In a linear system, it is possible to monitor the response of a series of arbitrary inputs to compute the inverse of the system’s transmission matrix (a mapping from inputs to outputs). This process entails measuring the responses of the whole system, which is practically not possible in a biological system given the large number of cells and the low number and resolution of the measuring electrodes. In practice, understanding the neural system occurs with partial measurements and it is a non-linear process. Thus, the development of an effective neural prosthesis lies in reducing the dimension of the external input while maximally preserving the information, so that few electrodes can write information in a format that the brain can read and understand.

To date, there have been several visual prostheses implanted in patients^17,26–31^, but most devices were tested to recognize letters and shapes using simple image encoding techniques (pixel averaging). In the Argus® II, the most implanted visual implant so far, pixel averaging in conjunction with video filters are used to down-sample the camera image to the resolution of the implanted array (6 x 10 pixels). Yet, there has been considerable research dedicated to the development of better image encoding algorithms. Some approaches include object detection, edge detection, and content-aware retargeting method^32,33^. In general, such methods aim to reduce the complexity of the image and highlight interesting content and features. For example, edge detection may identify the discontinuity of brightness in an image to locate the outline of an object. However, these algorithms do not always take into account the retinal information processing from photoreceptors to RGCs. Hence, they are not co-optimized with the elicited responses at the RGC level. Therefore, their encoding potential might be limited.

Concurrently, there had been significant efforts to generate in-silico retina models that potentially could be used for efficient image encoding^34–37^. Although, the literature on image encoding based on retinal modeling is scarce. Al-atabany et al. suggested a biophysical model that can maximize the useful visual information that can be transferred by simulating the different layers of retina^38^. Yet, the model used is critical since it will directly impact the outcome of the image encoding algorithm. Retinal information processing is complex^39,40^. In recent years, the use of convolutional neural networks (CNNs) have been very successful at modeling the retina and outperformed conventional approaches (linear-nonlinear model or generalized linear model)^41,42^. The latter have been shown to be less effective in capturing the retinal dynamics when natural scenes are presented^41^. CNNs have been shown to perform significantly better in modeling the retina for both white noise and natural scene stimuli^41,42^. Hence, using CNNs to model the retina presents a great potential in improving image encoding.

In this study, we propose an end-to-end neural network-based approach for both retina modeling and image encoding which takes into account the retinal information processing. We validate an actor-model framework designed to learn non-linear down-sampling patterns through a learning-based approach^43,44^. By integrating the measured retinal information processing into the framework, we demonstrate, in-silico and ex-vivo, that the generated down-sampled images elicit a neuronal response with higher neuronal reliability (+4.9% in-silico and +2.9% ex-vivo median percent increase) to the one produced by original images compared to a learning-free approach (pixel averaging). In addition, the artificial neural network model learned that contrast is a crucial feature for effective down-sampling.

The actor-model framework used in this study is general and can be exploited for other image encoding processes or even in other fields of sensory encoding, such as auditory and tactile. Albeit belonging to different sensory pathways, they share similar properties which allow us to postulate the potential effectiveness in other sensory encoding systems^40,45^. This methodological approach could serve as a template for future encoding strategies based on a learning-based approach to account for the natural transformation process occurring in the sensory organ. A more effective encoding method entails that the brain could better interpret the encoded information, leading to improved perception of a prosthesis user.

## RESULTS

### Actor-model framework in retinal processing

First, we conducted ex-vivo experiments to improve the image encoding step by accounting for the retinal information process. Using mouse retinal explants over a transparent multielectrode array (MEA), we recorded spiking activity from RGCs in response to projection of high-resolution images through the MEA onto the retinal photoreceptors (**Fig. 1a**). For each identified RGC, we built a response vector by summing up the number of spikes within a 400-ms window from the onset of each image. The image sequence was repeated 10 times to account for trial-to-trial variability, and responses to the same images were averaged.

**Figure 1.**
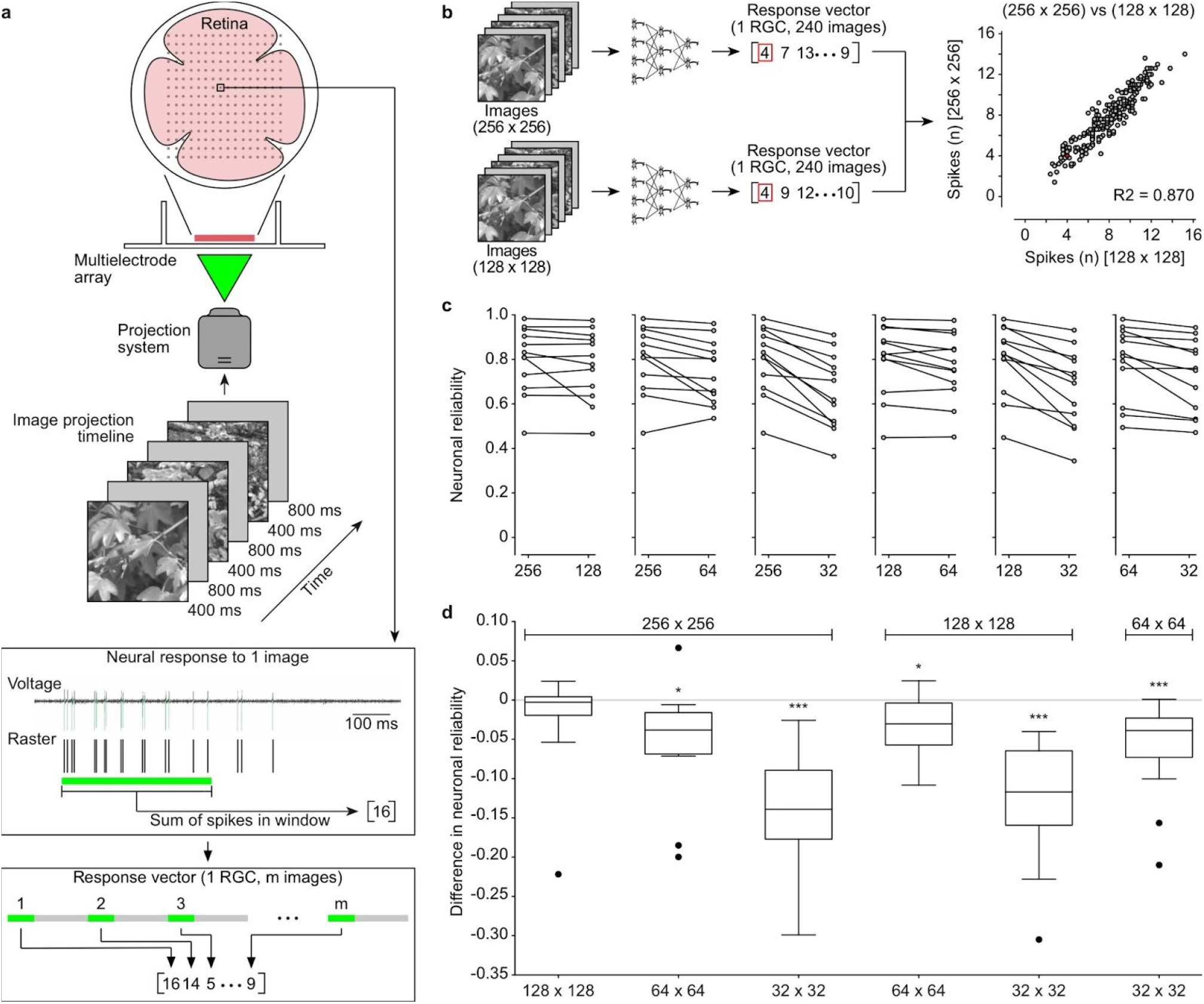
Experimental methodology. **a**, Images are projected onto the retina using a custom-made projection system. Each image is projected for 400 ms, and a gray frame is inserted between images for 800 ms to return the firing rate to baseline value. RGCs are isolated after spike detection and clustering using the SpyKING Circus algorithm. Action potentials from each recorded RGC from multiple explanted retinas are converted into raster plots. For each RGC, a response vector is built by summing up the number of spikes within a 400-ms window from the onset of each image. Responses to the repetition of the same image are averaged (not shown in the sketch) so numbers are not necessarily integers. The horizontal green bars correspond to the projected image. **b**, Representative example of neural reliability (R^2^) of one RGC comparing image dimension 256 x 256 pixels versus 128 x 128 pixels. The scatter plot is built from the two response vectors. The red circle is the data point corresponding to the two red boxes in the response vectors. **c**, The plots represent the paired distribution of neuronal reliabilities for all the recorded RGCs (n = 12 RGCs from N = 3 retinas). **d**, Each boxplot shows the distribution of the pairwise difference in neuronal reliability. The x-axis indicates the size of the low-resolution images while the high-resolution is indicated above the boxplot. The box represents the 25^th^ and the 75^th^ percentiles respectively, the line is the median, and the whiskers are 1.5 times the interquartile range. The black dots indicate outliers. p-values are reported as: * p < 0.05 and **** p < 0.0001.

In a pilot experiment, 240 images of different dimensions were projected (256 x 256, 128 x 128, 64 x 64, and 32 x 32 pixels) to determine the starting dimension of high-resolution images in subsequent experiments (**Fig. 1b**). Each block of 240 images was repeated 10 times. We recorded neural responses from 12 identified RGCs (from N = 3 retinas) and built 12 response vectors, for each of the four image resolutions. Then, we used neuronal reliability as a quantitative measure to compare the four resolutions in pairs. Briefly, the reliability of each RGC is measured as the R^2^ value between the neuronal responses to two paired images (**Fig. 1b**). Paired images are the same image presented at two different resolutions. Pooling all the RGCs together (**Fig. 1c**,**d**), we did not find any statistically significant difference in neuronal reliability when images of 256 x 256 pixels are down-sampled to 128 x 128 pixels by pixel averaging (p = 0.30, two-tailed paired Wilcoxon test). For all other comparisons, we found statistically significant differences (256 vs 64: p = 0.0122; 256 vs 32: p = 0.0005; 128 vs 64: p = 0.0200; 128 vs 32: p = 0.0005; 64 vs 32: p = 0.0009; two-tailed paired Wilcoxon tests). The choice of the resolution for high-resolution images is based on this result and other considerations. On the one hand, we aimed to minimize the high-resolution size to reduce the number of parameters of the CNN that will be trained as the forward model. Having more parameters will result in greater computational complexity and runs a larger risk of overfitting. Since we did not find a statistically significant difference while down-sampling from 256 x 256 to 128 x 128 pixels, we rejected the 256 x 256 pixel resolution. On the other hand, we still want to maximize the high-resolution size so that it can give a statistically significant difference in neuronal reliability compared to the down-sampled images, hence we chose 128 x 128 pixels as the resolution for the high-resolution images.

To build the actor-model network, first we repeated the ex-vivo experiment (n = 60 neurons from N = 10 retinas) using 1200 unique images of 128 x 128 pixels with 10 repeats for each image (**Fig. 2**, step 1). Hence, we identified the corresponding 60 neural response vectors and combined them into a neural response matrix as the ground truth for subsequent training phases of the framework. In the second phase (**Fig. 2**, step 2), we trained a forward model to act as a digital twin of the retina. We considered various models, and a CNN was chosen. To train the forward model, we pass high-resolution images (960 unique images) into the CNN as input, generating a prediction of the response vectors (predicted response matrix). We calculate the Poisson loss against the prediction and ground truth to update the forward model. We then conduct hyperparameter optimization with a random search. Once the forward model is trained, we fix its weights. Then, we prepended another CNN, the actor model, which learns to down-sample given images (**Fig. 2**, step 3). We input high-resolution images into the actor model (960 unique images), which reduces them to lower-resolution images of 32 x 32 pixels (four-fold down-sampling). The choice of four-fold down-sampling was also derived from the pilot experiment (**Fig. 1**). Since the aim of the actor network is to down-sample the images that can elicit a higher neuronal reliability, a sufficient reduction in the neuronal reliability between the high-resolution images and average down-sampled images is required to fully leverage on the potential of the actor-model framework. We found that four-fold down-sampling exhibited a greater reduction in neuronal reliability. The low-resolution images then pass through the forward model, generating a predicted response matrix. Similar to the training of the forward model, we compare the predicted response of the lower-resolution images against the ground truth by calculating the Poisson loss. The loss is then used to update the actor model. The forward model remained constant since it was fixed in step 2. As the actor model is updated, it learns to distill pertinent features to down-sample images while generating a neuronal response similar to high-resolution images.

**Figure 2.**
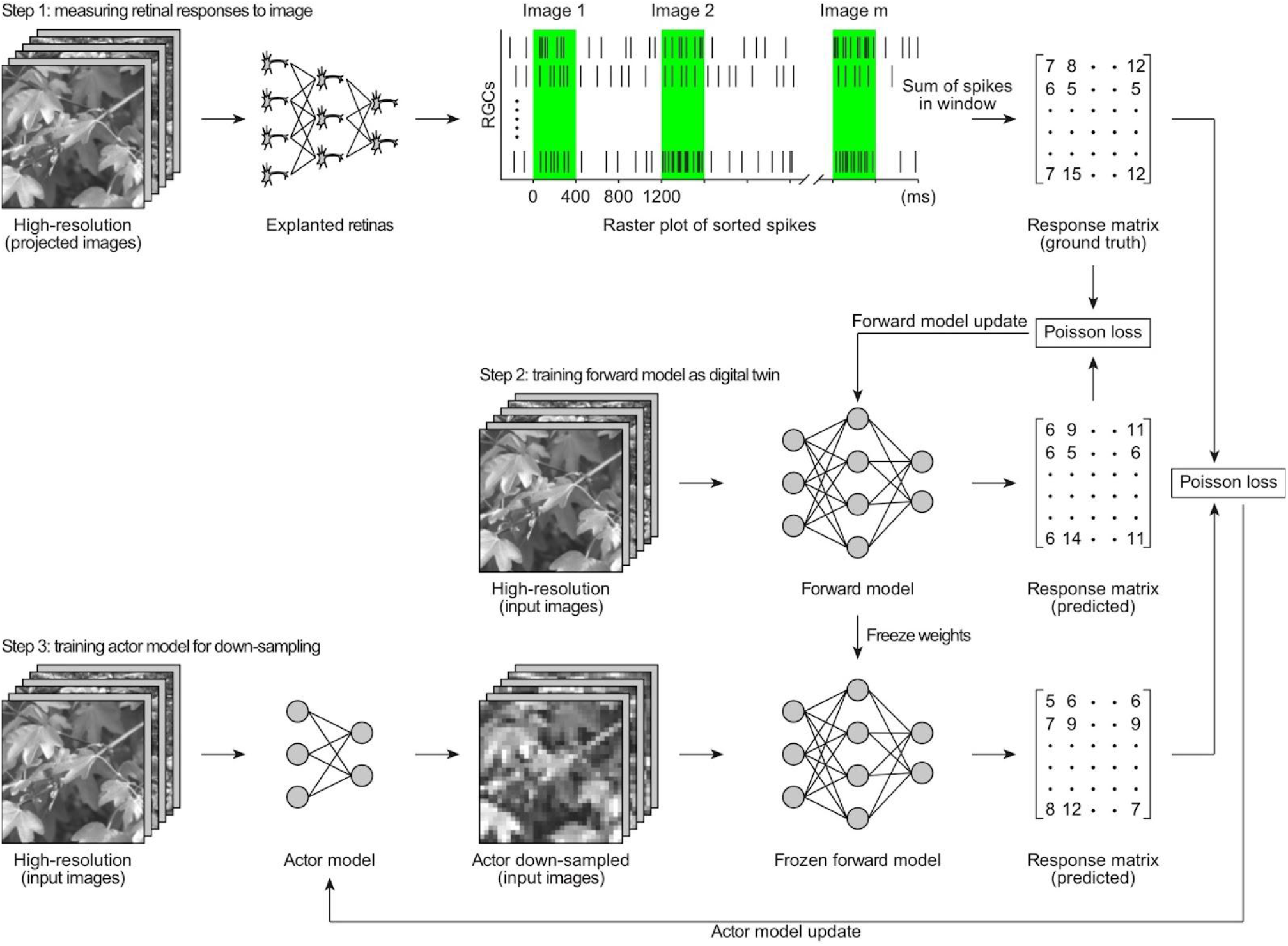
Actor-model framework for optimal downsampling. By projecting high-resolution images onto explanted retinas (green bars), we obtain the neuronal encoding of RGCs (Step 1). The raster plot is a representative sketch. This neuronal response matrix serves as the ground truth for the training of the forward CNN model. Given the same input, the predicted response of the model is compared against the ground truth by calculating the Poisson loss and used to update the forward model (Step 2). After training, the forward model is fixed and a CNN actor model is prepended. Similarly, high-resolution images are passed as input into the actor model which down-samples and feeds them into the fixed forward model. The predicted response is compared against the ground truth by calculating Poisson loss and used to update the actor model (Step 3).

### Actor-downsampled images elicit correlating responses to high-resolution images in-silico

With the forward and actor models trained, we conducted a comparison between the performance of the actor model and the pixel averaging method for four-fold down-sampling in-silico (**Fig. 3a**). Here, the forward model functions as the digital twin of an explanted mouse retina. We passed different types of images as inputs, including high-resolution images (high-resolution), images down-sampled by the actor network (actor down-sampled), and images down-sampled using the pixel averaging method (average down-sampled). For each group, we tested 120 unique images. Lastly, we computed the in-silico neuronal reliability. Briefly, the neuronal reliability of each RGC is measured as the R^2^ (**Fig. 3a**) between the neuronal responses of explanted retinas to high-resolution images (ground truth in **Fig. 2**) and the predicted responses of the forward model to the same high-resolution images, the average down-sampled images or the actor down-sampled images. The comparison between predicted response of high-resolution images and ground truth defines the baseline reliability of a digital RGC (**Fig. 3a**, magenta). The ground truth compared to predicted response of down-sampled images (**Fig. 3a**, actor in cyan or average in yellow) defines the in-silico neuronal reliability of down-sampled images, which is how similar the RGC responses to down-sampled images are to those from high-resolution images in-silico.

**Figure 3.**
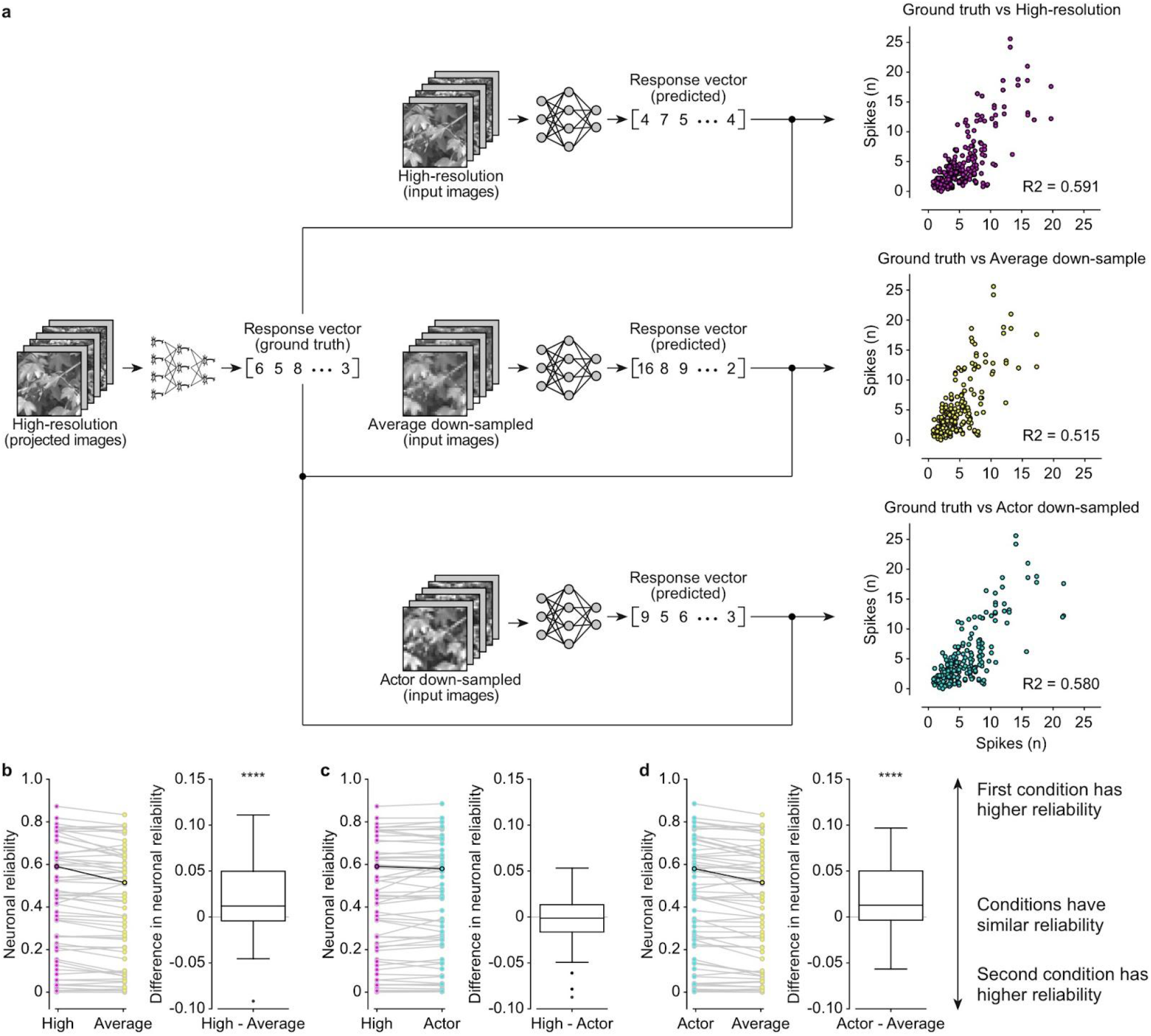
Results in-silico for four-fold downsampling. After training the forward and actor model, high-resolution, actor down-sampled and average down-sample images are passed as input to obtain the predicted response. **a**, For each neuron, we calculated the R-squared value of the predicted response and the ground-truth response. On the right is the scatter plot of the same representative neuron. **b-d**,The left plot represents the paired distribution of neuronal reliabilities for all the neurons (n = 60). Highlighted points correspond to the representative neuron in **a**. The boxplots show the distribution of the pairwise difference in neuronal reliability. The box represents the 25^th^ and the 75^th^ percentiles respectively, the line is the median, and the whiskers are 1.5 times the interquartile range. The black dots indicate outliers. p-values are reported as: **** p < 0.0001.

As expected, we observed a statistically significant reduction in neuronal reliability of the average down-sampled images compared to the baseline neuronal reliability (**Fig. 3b**; n = 60; p = 0.0001, two-tailed paired Wilcoxon test). However, we did not observe any statistically significant difference between the baseline neuronal reliability and the neuronal reliability of actor down-sampled images (**Fig 3c**; n = 60; p = 0.56, two-tailed paired Wilcoxon test). Furthermore, when comparing actor down-sampled images and average down-sampled images, the neuronal reliability of the actor-downsampled images was also significantly different (**Fig 3d**; n = 60, p = 0.0001, two-tailed paired Wilcoxon test). Although it appears that some neurons exhibit low in-silico neuronal reliability, this could be attributed to the learning of the model. The parameters for some of the neurons could be overfitted and since we do not model each neuron separately, it is difficult to ensure that every neuron is optimally modeled. Overall, we found a 4.9% median percent increase in neuronal reliability for actor down-sampled images compared to average down-sampled images. Based on these in-silico results, the actor network appears to have found a way that can elicit a neuronal response more similar to the ground truth compared to average down-sampling. The next logical step is to validate these in-silico findings ex-vivo in explanted retinas.

### Actor-downsampled images elicit correlating responses to high-resolution images ex-vivo

Finally, we validated the actor-model framework with a new set of retinal recordings. We explanted mouse retinas and measured the neuronal responses when presented with high-resolution images, actor down-sampled images, and average down-sampled images (n = 21 neurons from N = 8 retinas; 200 unique images, 10 repeats per image). High-resolution images were presented twice: first to determine a new ground truth, and then to compute neuronal reliability of high-resolution images. Then, we compared the performance of our learning-based down-sampling method with a learning-free downsampling method by calculating the ex-vivo neuronal reliability when compared to the new ground truth (**Fig. 4a**).

**Figure 4.**
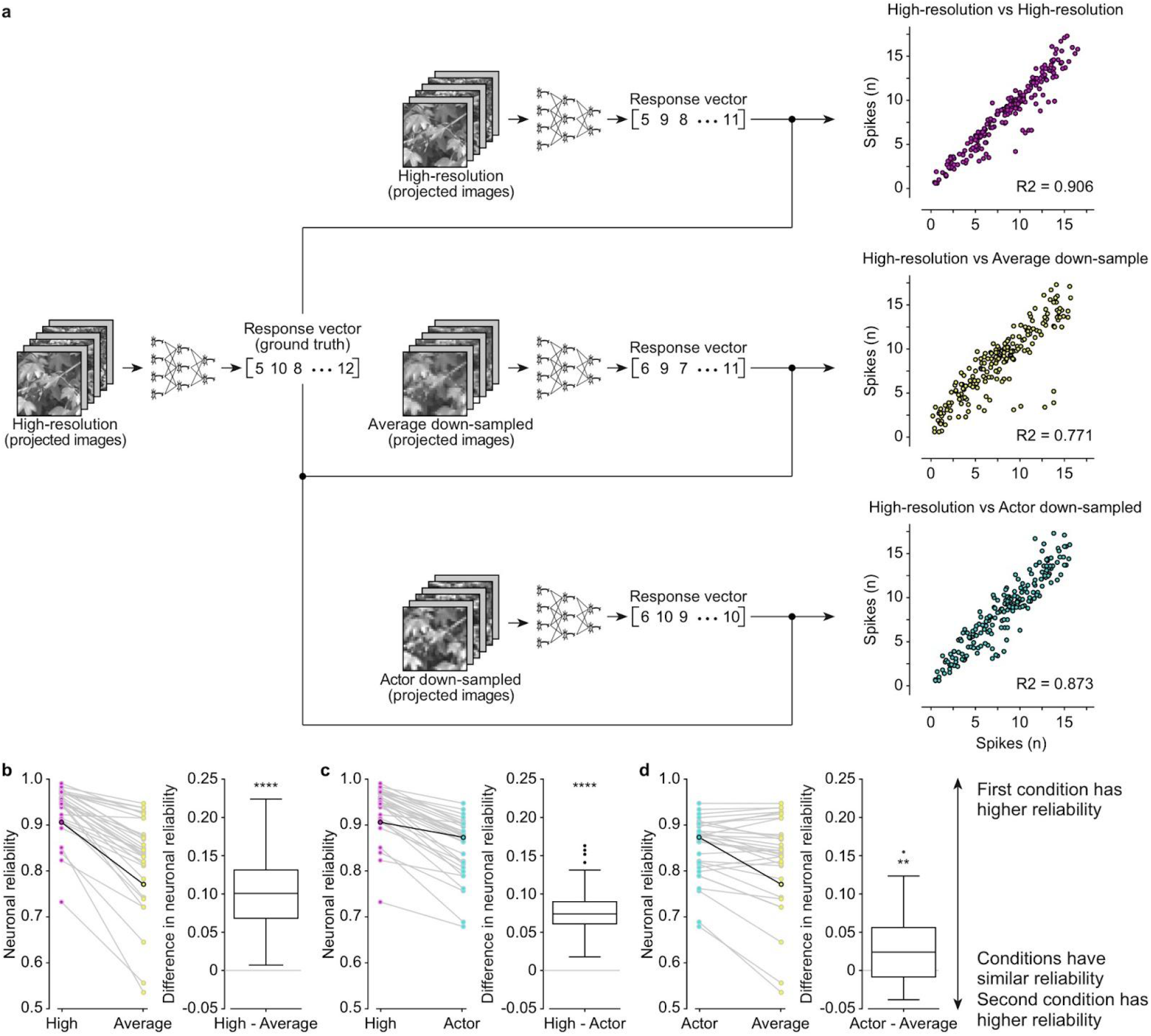
Results ex-vivo for four-fold downsampling. High-resolution, actor down-sampled and average down-sample images are projected onto the explanted retina over a MEA. **a**, For each neuron, we calculated the R-squared value of the neuronal response of high-resolution stimuli and paired stimuli. On the right is the scatter plot of the same representative neuron. **b-d**,The left plot represents the paired distribution of neuronal reliabilities for all the neurons (n = 21 from N = 8 retinas). Highlighted points correspond to the representative neuron in **a**. The boxplots show the distribution of the pairwise difference in neuronal reliability. The box represents the 25^th^ and the 75^th^ percentiles respectively, the line is the median, and the whiskers are 1.5 times the interquartile range. p-values are reported as: ** p < 0.01, and **** p < 0.0001

Qualitatively, results ex-vivo match in-silico data except for a significant difference between the baseline neuronal reliability and the neuronal reliability of actor down-sampled images (**Fig. 4**; n = 21, p < 0.0001, two-tailed paired Wilcoxon test). This was expected as information loss would occur during the down-sampling process. The baseline neuronal reliability is still significantly different from the neuronal reliability of average down-sampled images (**Fig. 4**; n = 21, p < 0.0001, two-tailed paired Wilcoxon test). Also, when comparing between actor down-sampled images and average down-sampled images, the neuronal reliability was significantly different (**Fig. 4d**; n = 21, p = 0.003, two-tailed paired Wilcoxon test). From the median percentage increase, we see that the actor down-sampling method performs about 2.9% better than the average down-sampling method.

### Actor down-sampling preserves contrasts of images

Then, we delved deeper into the pertinent attributes of effective down-sampling by comparing high-resolution images to their respective downsampled versions (**Fig. 5a**). Upon visual examination, it is evident that actor down-sampled images bear a greater resemblance to the original images. Furthermore, these images seem to exhibit a higher contrast in comparison to the average down-sampled images.

**Figure 5.**
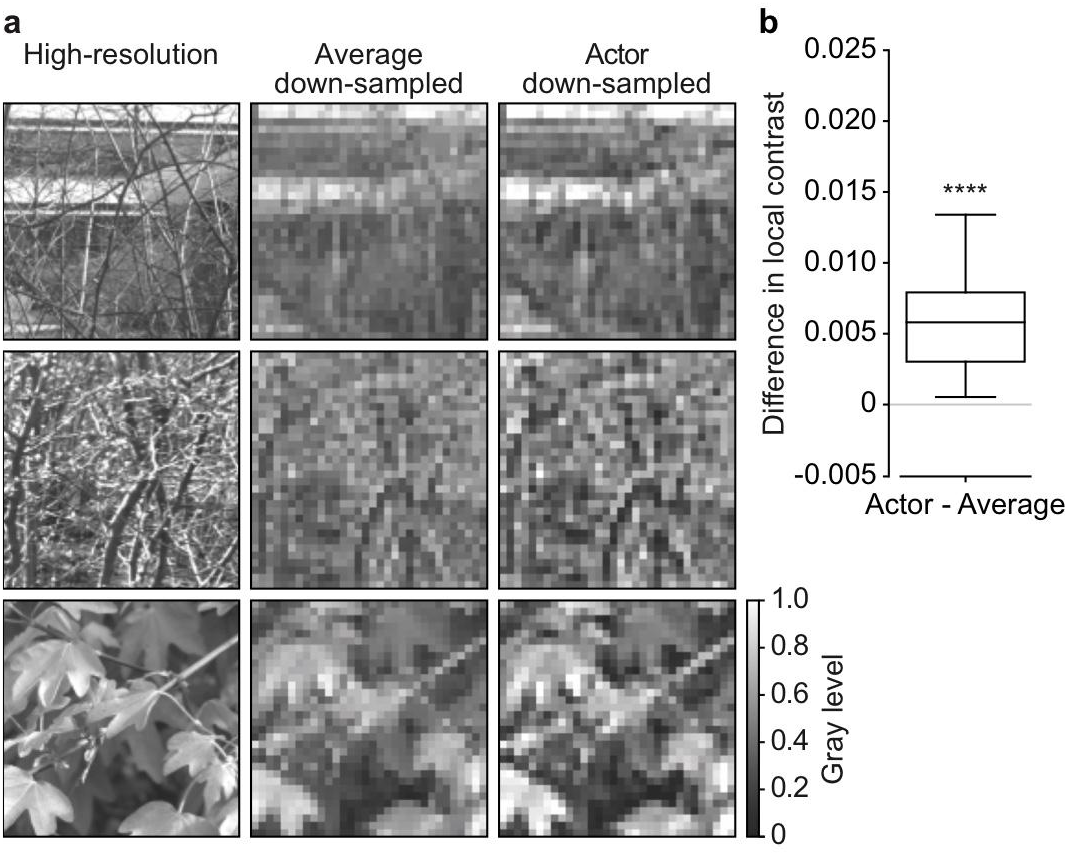
Results of different down-sampling methods. **a**. On the left, there are three high-resolution images with a resolution of 128 x 128 pixels. In the middle, the same images after four-fold average down-sampling. On the right, the same images after four-fold actor down-sampling. **b**. Distribution of the pairwise difference in local contrast between actor and average down-sampled images. The box represents the 25^th^ and the 75^th^ percentiles respectively, the line is the median, and the whiskers are 1.5 times the interquartile range. p-value is reported as: **** p < 0.0001.

This observation prompted us to quantify the local contrast of every down-sampled image by averaging the luminance variance of a sliding window of size 7 pixels x 7 pixels (**Fig. 5b**). The distribution of the local contrast difference shows that actor down-sampled images exhibit a significantly higher local contrast compared to average down-sampled images (p < 0.0001, one-tailed paired Wilcoxon test). This result implies that the actor model strives to increase the contrasts of the image, highlighting their significance in generating neuronal responses akin to those of high-resolution images.

### In-silico prediction of x-fold down-sampling

In the previous sections, we explored and validated the effectiveness of the actor-model framework during four-fold down-sampling, where images were reduced from 128 x 128 pixels to 32 x 32 pixels. The actor-model framework elicited a higher neuronal reliability (4.9% in-silico and 2.9% ex-vivo) compared to a learning-free approach (pixel averaging). Therefore, we decided to vary the folds of down-sampling in-silico to understand the extent of aggregation before the information loss cannot be recovered by the framework (**Fig. 6**). We trained multiple actor networks using similar method as described in the preceding sections (960 unique images) to down-sample the high-resolution images, each learning a different down-sampling fold: 2-fold (64 x 64 pixels), 4-fold (32 x 32 pixels), 8-fold (16 x 16 pixels), 16-fold (8 x 8 pixels), and 32-fold (4 x 4 pixels). We observe significant differences in neuronal reliability between actor down-sampled images and average down-sampled images up to 8 fold down-sampling (n = 60; 2 fold: p = 0.0178; 4 fold: p = 0.0004; 8 fold: p = 0.0006; two-tailed paired Wilcoxon tests). However, from 16 fold onwards, this difference was not observed (n = 60; 16 fold: p = 0.8024; 32 fold: p = 0.4483; two-tailed paired Wilcoxon tests). This was expected as 16 fold down-sampling corresponds to reducing the original image from 128 x 128 pixels to 8 x 8 pixels. Therefore, each pixel has a size of 400 x 400 μm^2^, exceeding the receptive field size of most RGC in the mouse retina^46^. Hence, the amount of information loss during down-sampling might be too much to be recovered.

**Figure 6.**
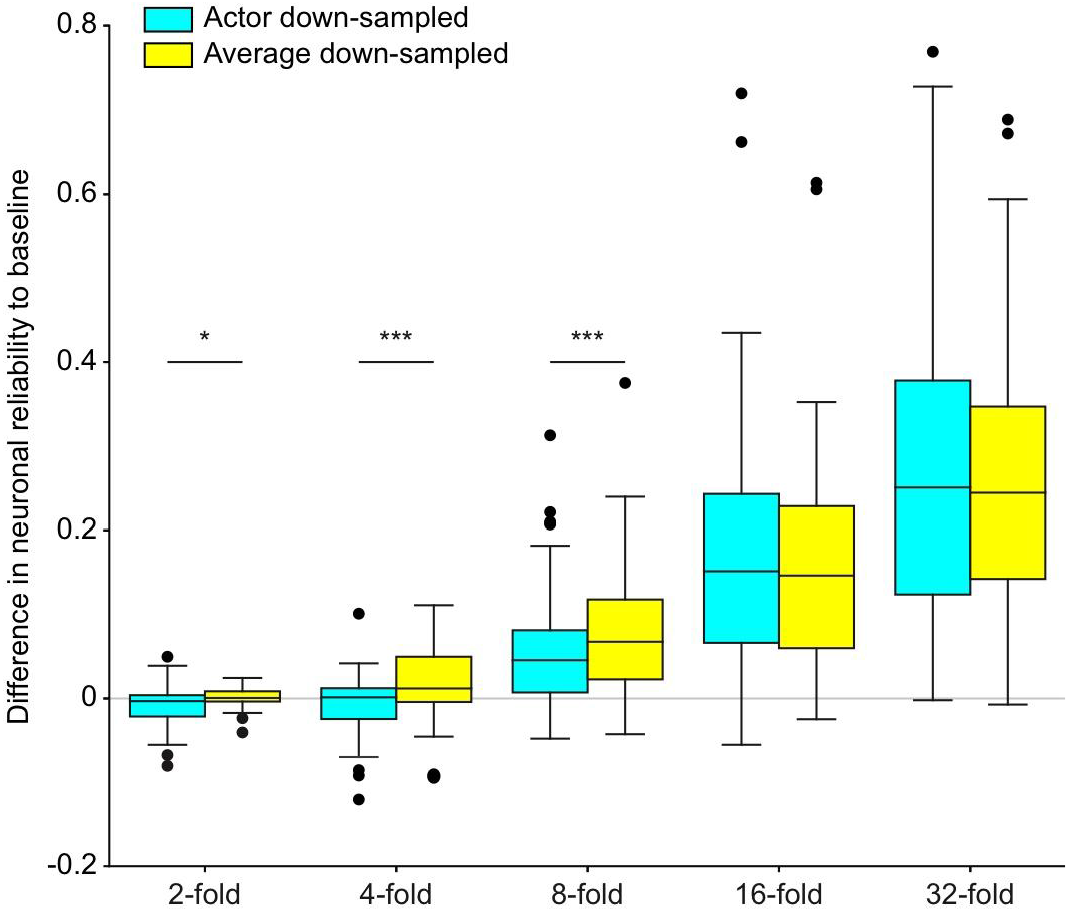
In-silico difference in neuronal reliability compared to baseline for x-fold down-sampling. Each boxplot is the distribution of the pairwise difference in in-silico neuronal reliability between the down-sampled images (actor down-sample in cyan; average down-sample in yellow) and high-resolution images for different down-sampling folds. The box represents the 25^th^ and the 75^th^ percentiles respectively, the line is the median, and the whiskers are 1.5 times the interquartile range. p-values are reported as: * p < 0.05, and *** p < 0.001.

Although we observed a significant difference between the neuronal reliability of actor down-sampled images and average down-sampled images for 2-fold down-sampling, a significant difference was not observed between neuronal reliability of average down-sampled images to the baseline (p = 0.1925, two-tailed paired Wilcoxon test), while actor down-sampled images has slightly higher neuronal reliability than baseline (p = 0.0277, one-tailed paired Wilcoxon test; p = 0.0139, one-tailed paired Wilcoxon test). This result suggests that the forward model is robust to down-sampling. This is not out of expectation as we used a CNN to model the retina, which could possess a certain level of robustness to down-sampling.

## DISCUSSION

In this study, we have successfully illustrated the efficacy of employing an actor-model framework to learn a down-sampling method that outperforms a standard learning-free pixel-average technique^27^. Furthermore, we substantiated the effectiveness of our proposed approach by analyzing the neuronal responses of RGCs. In this section, we delve into the implications of our findings and propose potential avenues for advancing the field of image encoding research.

Previously, the actor-model framework was employed within a physical system, where it successfully learned the necessary input to yield any desired output when transmitted through a multimode fiber. The performance achieved in that study was comparable to gold-standard methods^43^. In the present investigation, we sought to harness the potential of the actor-model framework within a biological system, which is often characterized by complexity and high variability.

In contrast to a physical system, we encountered a distinct set of challenges during the experimentation process. Firstly, like in many biological ex-vivo experiments, the difficulty lay in the sample viability, which constrained the duration of data collection. Given that neural networks typically require vast amounts of data to be effective, this limitation poses a significant obstacle to training the neural network efficiently. Secondly, the number of neurons recorded could fluctuate depending on the quality of the dissected tissue and other factors. Although we used 256 electrodes, not every electrode could successfully capture the electrical activity of a RGC. These issues imposed conducting experiments on multiple retinas and consolidating the recorded RGCs into a single dataset. As such, the framework has to learn and generalize across different mice to capture inter-sample variability, which could add another layer of difficulty. Lastly, intrinsic variability within the retina presented another hurdle. The response could vary from trial to trial when the same image was projected twice, with this variability being more pronounced than in the physical systems used in the previous work. Consequently, our actor-model framework required greater robustness to be effective.

Nevertheless, the actor-model framework successfully learned efficient down-sampling. This achievement underscores the robustness of the actor-model framework and its capacity to discover solutions irrespective of whether it is applied to a physical or biological system. We contend that the full potential of the actor-model has not yet been realized, and we look forward to future experiments that can capitalize on this versatile framework. For example in the context of tactile feedback, Eldeep and Akcakaya demonstrated the potential of using electroencephalography (EEG) to guide the electrical stimulation parameters^47^. The actor-model framework could also be applied in this situation, where the actor could determine the electrical stimulation required to elicit the desired EEG response.

By considering the biological neural network and constructing a digital twin of the retina using a CNN forward model, we successfully trained an actor network to down-sample images more efficiently than pixel averaging. In recent years, neural networks have excelled at modeling the retina, suggesting a level of complexity hidden within its seemingly simple structure^41,42^. As visual technologies progress and electrode resolution increases, image encoding methods become increasingly crucial. This study demonstrates that a learning-based approach, which accounts for biological retinal processes, yields superior results. We foresee this learning-based method serving as a foundation for future research, ultimately leading to the identification of the most effective image encoding technique. Additionally, this study hints at the importance of contrast during down-sampling, in agreement with a recent publication emphasizing the importance of contrast for a CNN model of the retina^41^. This result further highlights the benefits of utilizing neural networks for modeling to capture relevant underlying patterns. Employing a learning-based approach enables further analysis of learned features, providing insights into the underlying dynamics.

Additionally, this study shows the benefits of using CNNs for image encoding. The weight-sharing properties of CNNs allow for generalization across different retinas. As previously mentioned, a significant challenge in biological experiments involves conducting multiple trials with distinct samples. Neurons are recorded from multiple sessions and combined into a single dataset. However, even with such methods, not every area of the image is captured by the neuronal activity. Nevertheless, since weights are shared within a CNN, the actor network can extract pertinent features based solely on the subset of neurons measured. These distilled features can then generalize to other retinas and to other areas of the images not captured by neurons, as demonstrated by the positive results obtained in the ex-vivo experiments for validation.

Finally, it is important to note that our proposed actor down-sampling method is compatible with current image encoding techniques, such as saliency-based detection. These methods identify objects of interest to the user and encode only those objects. Our proposed method could complement such saliency-based detection^32,33,38,48^. First, we apply saliency detection algorithms to extract important objects, and then pass the processed images through the actor model for more effective downsampling.

As a preliminary step toward a learning-based approach for image encoding, there remains ample room for improvement and exploration. In this study, we projected static images and summed the number of spikes within a specified window. Constrained by the hardware employed, we were unable to present images in a continuous format (i.e., movie format), which prevented us from verifying whether our proposed methods would be applicable to more dynamic natural scenes. A logical next step would be to validate these results using continuous projections of natural scenes. In addition to exploring continuous format, the same approach could be evaluated on reducing the bit size of the depth of the images concurrently while down-sampling. Reducing from 256 levels of grayscale to 8 levels would also be useful for visual prostheses to better calibrate the strength of electrical stimulation.

Another avenue is investigating the generalization of our approach across species. The actor and forward models were trained using data collected from mouse retinas. We also discovered that the results derived from these trained models could be generalized across different mouse retinas, as the validation experiments were conducted with new sets of retinas. It would be interesting to explore whether projecting the various down-sampled images onto the retinas of another species would yield similar improvements. If successful, this would imply that the features learned by the actor model may possess the capacity to generalize even across different species, highlighting the potential for broader applicability of this method.

In conclusion, this study presents a novel neural network-based approach for optimizing image distillation in the context of visual prostheses. The proposed actor-model framework learns to downsample images while accounting for the biological processes of the retina, resulting in more effective down-sampling patterns. This research not only contributes to the advancement of image encoding techniques for visual prostheses but also highlights the importance of incorporating natural biological transformations to improve the perceptual experience of prosthetic users. Future research could build upon this learning-based approach to develop even more accurate and effective image encoding methods, ultimately enhancing the quality of life for individuals relying on such devices.

## METHODS

### Electrophysiological recordings

Animal experiments were authorized by the Direction Générale de la Santé de la République et Canton de Genève in Switzerland (authorization number GE31/20). C57BL/6J mice (n = 21; age 75.3 ± 26.4 days, mean ± SD) were dark-adapted for 1 hr prior to euthanasia. Euthanasia was carried out via intraperitoneal injection of sodium pentobarbital (150 mg kg^−1^). Dissection and recording of retinas were performed in carboxygenated (95% O_2_ and 5% CO_2_) Ames medium (USBiological, A1372-25) under dim red light. Retinas were maintained at 25°C throughout the experiment. Explanted retinas were positioned on a poly-lysine coated membrane (Sigma, P8920; Repligen 132544), with the RGCs side facing a 256-channel MEA (256MEA200/30iR-ITO, Multichannel systems) with 30-μm electrodes spaced 200 μm. The data sampling rate was set at 25 kHz.

### Visual stimulation

Images were projected using a custom-built setup with a Digital Mirror Device (V-7000 Hi-Speed V-Modules, ViALUX) coupled to a white LED (MWWHF2, Thorlabs). The stimulus was focused on the photoreceptors via standard optics, with an average power of 13 nW. The projected area covered 3.2 x 3.2 mm^2^. The image set is the van Hateran Natural Image Dataset, comprising 4212 monochromatic and calibrated images captured in a variety of natural environments^49^, further processed to maintain a linear relationship between scene luminance and pixel values^41^. This processing step was described as crucial to prevent the retinal system from having to adapt to varying light intensity levels found in different environments^41^. The final image database used in this experiment (3190 images) was obtained from Goldin et al.^41^, which was then sub-sampled for the different experiments conducted.

### Spike detection and sorting

The SpyKING CIRCUS algorithm was used for spike sorting^50^. Manual inspection was performed using Phy software^51^. To filter out poor-quality clusters, we presented a random binary checkerboard at the end of the experiment for 1 hr at 33 Hz^52^. The check size was 50 μm. The receptive field of each spike-sorted cluster was mapped. Clusters that did not display a clear receptive field were considered non-neuronal and excluded from further analysis. To assess the reliability of the recorded neurons and account for experimental drift, an identical set of binary checkerboard stimuli was initially displayed at the start of the experiment, and then redisplayed roughly every half an hour. The correlation coefficient of a cell’s average response to the same stimulus across different blocks of trials was calculated. Only neurons with a correlation exceeding 0.3 were selected for further analysis^42^.

### Model architecture

The forward model architecture adheres to the current state of the art^41,53,54^. It consists of two layers, the first layer of the CNN, *k*_*rsk*_, convolutes the input image. The output of the first convolution layer passes through a pointwise nonlinear function, 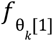, to obtain non-negative activation values. For each neuron n, a readout weight *w*_*ijkn*_, which factorizes as *w*_*ijkn*_ = *u*_*ijn*_ *ν*_*kn*_, is applied. Here, *i* and *j* index space, with *u*_*ijn*_ representing spatial weights and *ν*_*kn*_ denoting feature weights. Another nonlinear activation function, 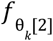, is performed, followed by the utilization of a Poisson noise model during training. Softplus was chosen as the activation function for 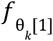 and 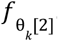. Essentially, the k^th^ neuron of the first layer is represented as 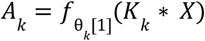. The spiking rate *r*_*n*_ of the n^th^ neuron given an input image *X* was 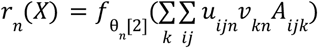. Additionally, batch normalization was applied to the outputs of the first layer. Laplacian regularization was applied to the convolutional kernels of the first layer. For the feature and spatial weights of the second layer, we used L1 regularization such that: 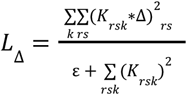 and 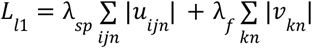 with ε = 10^−8^. The actor model consists of a convolution kernel that was prepended to the forward model (**Supplementary Fig. 1**).

### Neural networks training

Given n image-response pairs (*X*_1_, *y*_1_), …, (*X*_*n*_, *y*_*n*_), the loss function of the forward network is provided by: 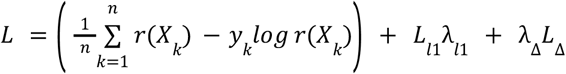. The first term corresponds to the Poisson loss, while the second and third terms represent the regularization terms. The model was fitted with the Adam optimizer on the training set. The responses were obtained by summing up the number of events elicited within 400 ms from the image onset. We maintained a constant batch size of 32 during training for both actor and forward networks. For the learning rate, we started with 0.001 for forward network and 0.002 for actor network. To avoid overfitting, we employed both early stopping and the decay mechanism with maximum 1000 epochs (**Supplementary Fig. 2**). The hyperparameters for the regularization term were optimized by performing a random search for the forward network, while a grid search was employed for the actor network. The optimal hyperparameter values were the ones whose model produced the lowest loss value without regularization terms on the validation dataset. The hyperparameters for the forward network were 0.0033 for smoothing factor of convolution kernels, 0.00278 for spatial sparsity factor and 1.34^-6^ for feature sparsity factor. For the actor network, the best run had 6 convolution kernels of size 31 x 31 with 0.1 L2 regularization.

### Neuronal reliability

As a quantitative measurement of performance for both in-silico and ex-vivo experiments, we calculated the R^2^ value. For each neuron, we calculated the R^2^ value for each input image and found the average across the images and neurons, such that: 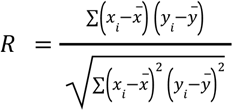. Where x and y can be the predicted response or actual neuronal response to different sets of images

### Statistical analysis

Statistical analyses were conducted with the Python Scipy library (python version 3.10.6, scipy version 1.9.1). The Shapiro-Wilk normality test was performed to justify the use of non-parametric tests.

## DATA AVAILABILITY

The authors declare that the data supporting the findings of this study are available in the paper. The source data file is provided with this manuscript.

## CODE AVAILABILITY

The code used in this study is available at https://github.com/lne-lab/actor-retina

## ACKNOWLEDGEMENTS

We would like to thank Silvestro Micera (EPFL, Switzerland) for reading the manuscript and providing valuable feedback. We would also like to thank the team of Olivier Marre (Institut de la Vision, France) for providing the code and preprocessed images used in this manuscript. The project was supported by the EPFL STI e-seed fund.

## SUPPLEMENTARY FIGURES

**Supplementary Fig. 1.**
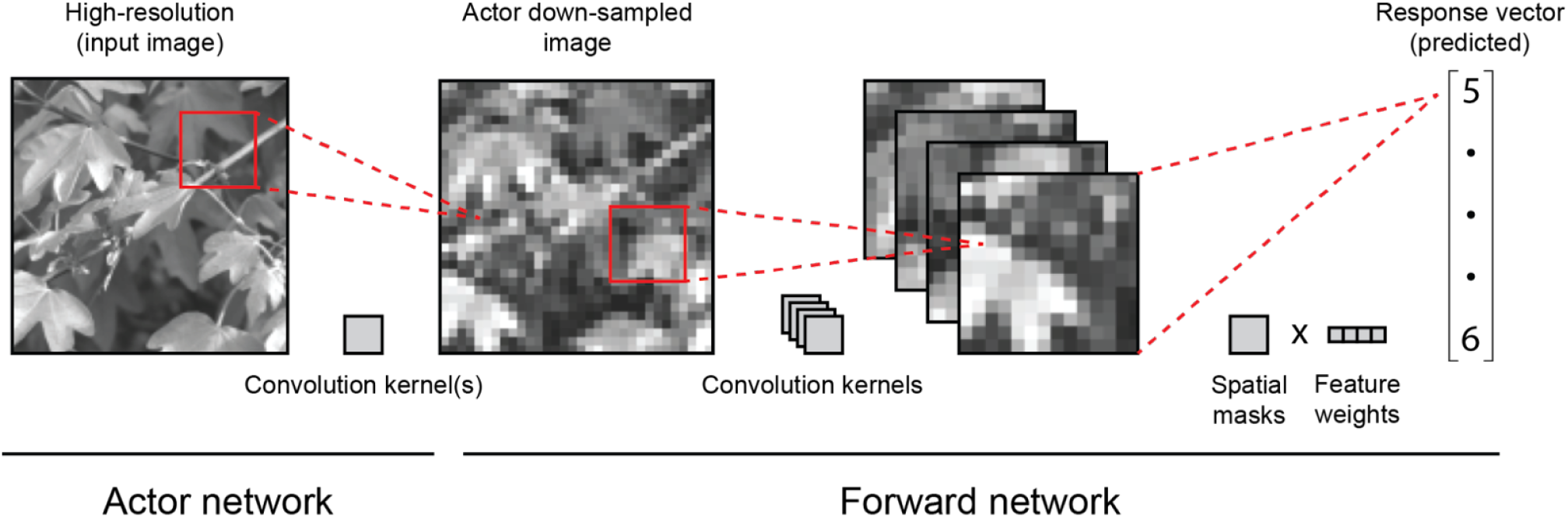
Network architecture. The forward network comprises a layer of convolution kernels, followed by the readout layer, which is decomposed into spatial masks and feature weights. Prepended to the trained forward network is the actor network, which consists of a layer of convolution kernels. The schematic illustrates the flow as we pass a single image as input through the framework to generate the predicted response vector of the neurons.

**Supplementary Fig. 2.**
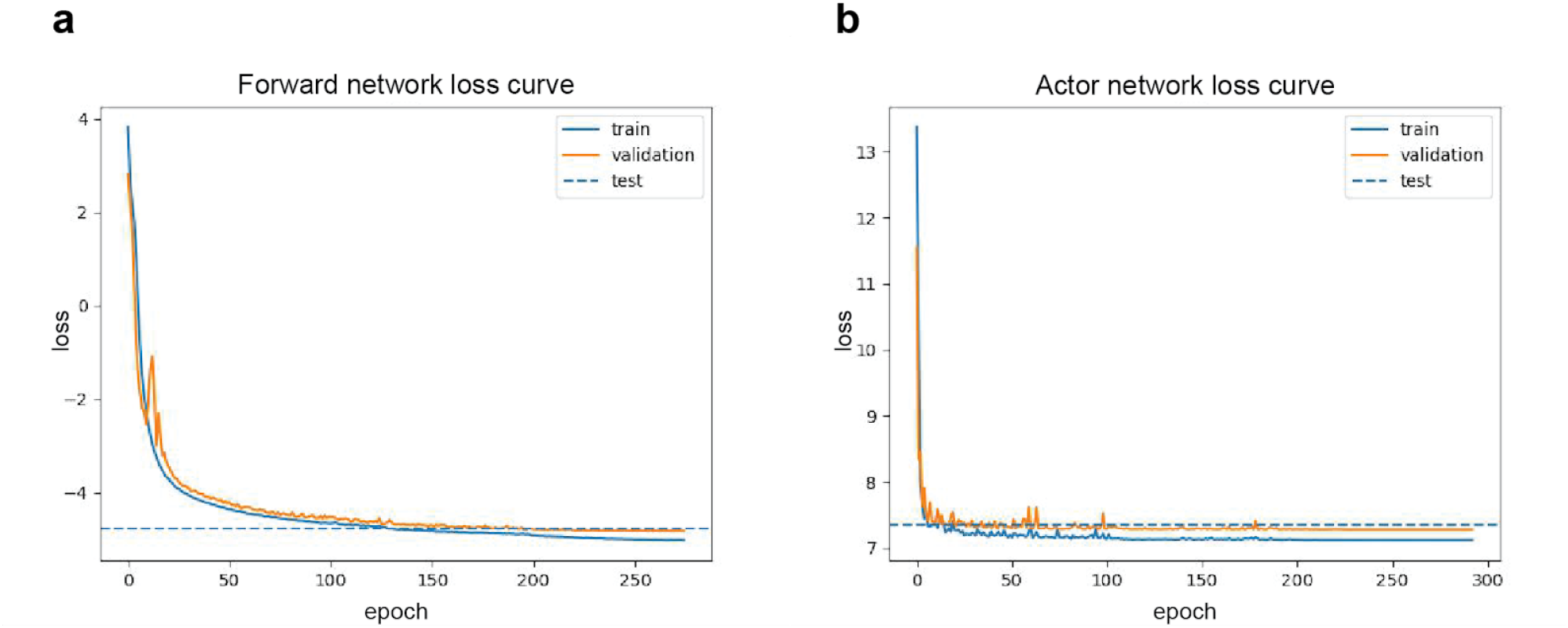
Training curve of the networks. **a**. Training loss of forward network. **b**. Training loss of actor network. The total loss of the training was plotted against the epoch. We observe a steady decrease in the training and validation loss without signs of divergence. The loss value of the test dataset was also similar to the training and validation.

## BIBLIOGRAPHY

1. Wilson, B. S. & Dorman, M. F. Cochlear implants: A remarkable past and a brilliant future. Hear. Res. 242, 3–21 (2008).

2. Wilson, B. S. et al. Better speech recognition with cochlear implants. Nature 352, 236–238 (1991).

3. Zeng, F.-G., Rebscher, S., Harrison, W., Sun, X. & Feng, H. Cochlear Implants: System Design, Integration, and Evaluation. IEEE Rev. Biomed. Eng. 1, 115–142 (2008).

4. George, J. A. et al. Biomimetic sensory feedback through peripheral nerve stimulation improves dexterous use of a bionic hand. Sci. Robot. 4, p(2019).

5. Raspopovic, S. et al. Restoring Natural Sensory Feedback in Real-Time Bidirectional Hand Prostheses. Sci Transl Med 6, 222ra19 (2014).

6. Tabot, G. A. et al. Restoring the sense of touch with a prosthetic hand through a brain interface. Proc. Natl. Acad. Sci. 110, 18279–18284 (2013).

7. Dhillon, G. S. & Horch, K. W. Direct Neural Sensory Feedback and Control of a Prosthetic Arm. IEEE Trans. Neural Syst. Rehabilitation Eng. 13, 468–472 (2005).

8. Tan, D. W. et al. A neural interface provides long-term stable natural touch perception. Sci. Transl. Med. 6, 257ra138 (2014).

9. Osborn, L. E. et al. Prosthesis with neuromorphic multilayered e-dermis perceives touch and pain. Sci. Robot. 3, (2018).

10. Bensmaia, S. J., Tyler, D. J. & Micera, S. Restoration of sensory information via bionic hands. Nat. Biomed. Eng. 7, 443–455 (2023).

11. Losanno, E., Mender, M., Chestek, C., Shokur, S. & Micera, S. Neurotechnologies to restore hand functions. Nat. Rev. Bioeng. 1, 390–407 (2023).

12. Vagni, P. et al. POLYRETINA restores light responses in vivo in blind Göttingen minipigs. Nat Commun 13, 3678 (2022).

13. Chenais, N. A. L., Leccardi, M. J. I. A. & Ghezzi, D. Photovoltaic retinal prosthesis restores high-resolution responses to single-pixel stimulation in blind retinas. Commun Mater 2, 28 (2021).

14. Gaillet, V. et al. Spatially selective activation of the visual cortex via intraneural stimulation of the optic nerve. Nat Biomed Eng 4, 181–194 (2020).

15. Gaillet, V., Borda, E., Zollinger, E. G. & Ghezzi, D. A machine-learning algorithm correctly classifies cortical evoked potentials from both visual stimulation and electrical stimulation of the optic nerve. J Neural Eng 18, 046031 (2021).

16. Chen, X., Wang, F., Fernandez, E. & Roelfsema, P. R. Shape perception via a high-channel-count neuroprosthesis in monkey visual cortex. Science 370, 1191–1196 (2020).

17. Fernández, E. et al. Visual percepts evoked with an Intracortical 96-channel microelectrode array inserted in human occipital cortex. J Clin Invest 131, p(2021).

18. Jeon, C.-J., Strettoi, E. & Masland, R. H. The Major Cell Populations of the Mouse Retina. J Neurosci 18, 8936–8946 (1998).

19. Masland, R. H. The Neuronal Organization of the Retina. Neuron 76, 266–280 (2012).

20. Masland, R. H. Accurate maps of visual circuitry. Nature 500, 154–155 (2013).

21. Masland, R. H. Neuronal diversity in the retina. Curr Opin Neurobiol 11, 431–436 (2001).

22. Valle, G. et al. Biomimetic Intraneural Sensory Feedback Enhances Sensation Naturalness, Tactile Sensitivity, and Manual Dexterity in a Bidirectional Prosthesis. Neuron 100, 37–45.e7 (2018).

23. Mendez, V., Iberite, F., Shokur, S. & Micera, S. Current Solutions and Future Trends for Robotic Prosthetic Hands. Annu. Rev. Control, Robot., Auton. Syst. 4, 1–33 (2021).

24. Wang, B.-Y. et al. Electronic photoreceptors enable prosthetic visual acuity matching the natural resolution in rats. Nat Commun 13, 6627 (2022).

25. Borda, E. & Ghezzi, D. Advances in visual prostheses: engineering and biological challenges. Prog Biomed Eng 4, 032003 (2022).

26. Luo, Y. H.-L. & Cruz, L. da. The Argus® II Retinal Prosthesis System. Prog Retin Eye Res 50, 89–107 (2016).

27. Edwards, T. L. et al. Assessment of the Electronic Retinal Implant Alpha AMS in Restoring Vision to Blind Patients with End-Stage Retinitis Pigmentosa. Ophthalmology 125, 432–443 (2018).

28. Palanker, D., Mer, Y. L., Mohand-Said, S. & Sahel, J. A. Simultaneous perception of prosthetic and natural vision in AMD patients. Nat Commun 13, 513 (2022).

29. Muqit, M. M. K. et al. Six-Month Safety and Efficacy of the Intelligent Retinal Implant System II Device in Retinitis Pigmentosa.Ophthalmology 126, 637–639 (2019).

30. Petoe, M. A. et al. A Second-Generation (44-Channel) Suprachoroidal Retinal Prosthesis: Interim Clinical Trial Results. Transl Vis Sci Technology 10, 12 (2021).

31. Veraart, C., Wanet-Defalque, M., Gérard, B., Vanlierde, A. & Delbeke, J. Pattern Recognition with the Optic Nerve Visual Prosthesis. Artif Organs 27, 996–1004 (2003).

32. Wang, J. et al. The application of computer vision to visual prosthesis. Artif Organs 45, 1141–1154 (2021).

33. Sanchez-Garcia, M., Martinez-Cantin, R. & Guerrero, J. J. Semantic and structural image segmentation for prosthetic vision. PLoS ONE 15, e0227677 (2020).

34. Beyeler, M., Boynton, G., Fine, I. & Rokem, A. pulse2percept: A Python-based simulation framework for bionic vision. Proc. 16th Python Sci. Conf. 81–88 (2017) doi:10.25080/shinma-7f4c6e7-00c.

35. Lozano, A., Garrigós, J., Martínez, J. J., Ferrández, J. M. & Fernández, E. Natural and Artificial Computation for Biomedicine and Neuroscience, International Work-Conference on the Interplay Between Natural and Artificial Computation, IWINAC 2017, Corunna, Spain, June 19-23, 2017, pProceedings, Part I. Lect. Notes Comput. Sci. 464–472 (2017) doi:10.1007/978-3-319-59740-9_46.

36. Lozano, A. et al. Neurolight: A Deep Learning Neural Interface for Cortical Visual Prostheses. Int J Neural Syst 30, 2050045 (2020).

37. Wienbar, S. & Schwartz, G. W. The dynamic receptive fields of retinal ganglion cells. Prog. Retin. Eye Res. 67, 102–117 (2018).

38. Al-Atabany, W., McGovern, B., Mehran, K., Berlinguer-Palmini, R. & Degenaar, P. A Processing Platform for Optoelectronic/Optogenetic Retinal Prosthesis. IEEE Trans. Biomed. Eng. 60, 781–791 (2013).

39. Field, G. D. & Chichilnisky, E. J. Information Processing in the Primate Retina: Circuitry and Coding. Annu. Rev. Neurosci. 30, 1–30 (2007).

40. Saal, H. P., Harvey, M. A. & Bensmaia, S. J. Rate and timing of cortical responses driven by separate sensory channels. eLife 4, e10450 (2015).

41. Goldin, M. A. et al. Context-dependent selectivity to natural images in the retina. Nat. Commun. 13, 5556 (2022).

42. McIntosh, L. T., Maheswaranathan, N., Nayebi, A., Ganguli, S. & Baccus, S. A. Deep Learning Models of the Retinal Response to Natural Scenes. Adv. neural Inf. Process. Syst. 29, 1369–1377 (2016).

43. Rahmani, B. et al. Actor neural networks for the robust control of partially measured nonlinear systems showcased for image propagation through diffuse media. Nat Mach Intell 2, 403–410 (2020).

44. Rahmani, B., Psaltis, D. & Moser, C. Variational framework for partially-measured physical system control: examples of vision neuroscience and optical random media. arXiv (2021) doi:10.48550/arxiv.2110.13228.

45. Pei, Y. C., Hsiao, S. S. & Bensmaia, S. J. The tactile integration of local motion cues is analogous to its visual counterpart. Proc. Natl. Acad. Sci. 105, 8130–8135 (2008).

46. Ran, Y. et al. Type-specific dendritic integration in mouse retinal ganglion cells. Nat. Commun. 11, 2101 (2020).

47. Eldeeb, S. & Akcakaya, M. EEG guided electrical stimulation parameters generation from texture force profiles. J. Neural Eng. 19, 066042 (2022).

48. Kasowski, J., Wu, N. & Beyeler, M. Towards Immersive Virtual Reality Simulations of Bionic Vision. Augment. Hum. Conf. 2021 313–315 (2021) doi:10.1145/3458709.3459003.

49. Hateren, J. H. van & Schaaf, A. van der. Independent component filters of natural images compared with simple cells in primary visual cortex. Proc. R. Soc. Lond. Ser. B: Biol. Sci. 265, 359–366 (1998).

50. Yger, P. et al. A spike sorting toolbox for up to thousands of electrodes validated with ground truth recordings in vitro and in vivo. eLife 7, e34518 (2018).

51. Rossant, C. et al. Spike sorting for large, dense electrode arrays. Nat. Neurosci. 19, 634–641 (2016).

52. Chichilnisky, E. J. A simple white noise analysis of neuronal light responses. Netw Comput Neural Syst 12, 199–213 (2001).

53. Klindt, D. A., Ecker, A. S., Euler, T. & Bethge, M. Neural system identification for large populations separating “what” and “where.” arXiv (2017) doi:10.48550/arxiv.1711.02653.

54. Cadena, S. A. et al. Deep convolutional models improve predictions of macaque V1 responses to natural images. PLoS Comput. Biol. 15, e1006897 (2019).

